# Optimizing PCR using mtDNA Universal Primers for Detecting Indigenous Domestic Animal Species from Pakistan

**DOI:** 10.1101/2023.12.02.569703

**Authors:** Hafiza Iqra Saeed, Mashmoom Suhail, Sehreem Gill, Rashid Saif

## Abstract

The prevalence of specific animal meat consumption is integral to human nutrition but is jeopardized by the widespread issue of meat adulteration, posing threats to both the industry and consumer health. Escalating demand has led to fraudulent practices, necessitating reliable techniques for authentic source identification, aligning with ethical and religious principles. PCR has proven effective in addressing this concern by examining twelve domestic animal species in Pakistan, differentiating between halal and haram species. The method employs universal primers for 16S rRNA gene amplification, generating species-specific DNA fragments. This sophisticated approach, producing distinct fragments for cow, camel, goat, sheep, chicken, cat, dog, donkey, duck, rabbit, horse, and human with 507, 385, 787, 763, 470, 700, 370, 600, 500, 750, 800 and 300bp holds promise for identifying and tracing these species in commercial meat products. By ensuring robust food traceability, this methodology combats contemporary challenges in adulteration, preserving the integrity of the food supply chain.

## Introduction

Consumption of meat increases with gradual increase in population. Mislabelling and adulteration of meat products have been escalated previously. Consumers demand quality products with complete nutritional attributes in low prices but increased fraudulency make it impossible. Meat is the richest source of protein and provides essential nutrients. The people engaged in fraudulency with the intention of making more profit at cheaper rate, either substitutes the originally demanded specie with a low quality or entirely different specie or a mixture of species being common possibilities [2]. This fraudulent act of contamination breaks or violates ethical laws of food safety [3] as Hindus and Jews do not eat Cow’ meat and Muslims are prohibited from haram meat (Donkey, dog, turtle, pork etc.). Pakistan is facing drastic peak in meat adulteration cases in past few years. The scenario clearly projects non-compliance of halal food laws [4]. This concrns the public health as sometimes edible and inedible meat can elicit the allergies, metabolic disorders and infectious diseases specially in sensitive and immunocompromised individuals [5].

To protect the consumers being deceived and to ensure the safety of dietary products; molecular detection of meat is very important. Numerous analytical techniques have been used to detect the meat and meat products. Protein based tecniques such as ELISA is not fully compatible as proteins gets denatured at extreme temperatures. For this purpose molecular DNA based techniques are cost effective and provide accurate results. PCR based techniques have become the key to finding fraudulent components in raw and processed meat [6].

In the current study, a set of universal primers has been used for mitochondrial 16S rRNA against twelve domestic species from Pakistan. *Bos taurus* (cow), *Camelus bactrianus* (camel), *Capra hircus* (goat), *Ovis aries* (sheep), *Gallus domesticus* (chicken), *Felis catus* (cat), *Canis familiasris* (dog), *Equus asinus* (donkey), *Anas Platyrhynchos* (duck), *Oryctolagus cuniculus* (rabbit), *Equus caballus* (horse), and *Homo sapiens* (human). The objective of this study is to focus on finding whether the pair of universal primers amplify the mitochondrial target region in the recruited animal species or not allowing the identification of species in their meat products.

## Material and Methods

### Sample collection and DNA Extraction

Blood samples were collected from twelve different animal species, local to Pakistan. The samples were collected in K3-EDTA tubes and stored at 4°C for further processing. DNA was extracted from each sample using GDSBio genomic DNA extraction kit.

### Mitochondrial 16s rRNA Universal Primers

A set of universal primers were taken for the identification of aforementioned twelve animal species for their mitochondrial DNA (mtDNA) amplification [7]. Forward and reverse primers were blast against all the sample species using an online tool in the NCBI database https://www.ncbi.nlm.nih.gov/tools/primer-blast/. The available sequences of each species showed positive results with the set of universal primers, making species identification possible before going into the wet-lab protocols.

**Table 1:**
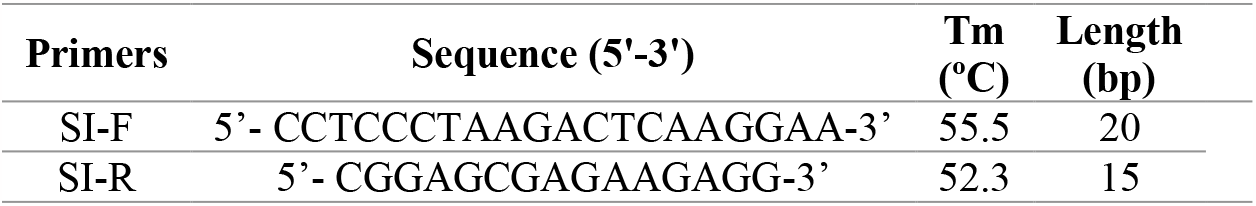
Sequences and attributes of PCR primers used for species identification.

### PCR Amplification

PCR reactions were carried out on a SimpliAmp-applied biosystems thermal cycler with 20µL of total reaction mixture containing 200mM dNTPs, 2.0µL of 10X PCR buffer, 1.8mM Mgcl_2_, 1IU of Taq polymerase, 100pmol of each universal primer, and 50ng/µL of DNA template. Cycling parameters for standard PCR were: preheating at 94°C for 5 min; 30 cycles of denaturation at 94 °C for 30sec, annealing at 55-60 °C for 30sec, and extension at 72 °C for 30sec; final extension at 72 °C for 5 min. A 5.0µL of PCR product was analyzed by 1% agarose gel electrophoresis (about 35 min at 80V) in 1X TAE buffer.

## Results

PCR products amplified from collected samples of twelve different species shown in figure 2. Each species is amplified with the same set of universal primers, which shows amplified mitochondrial target gene 16s rRNA that can act as a marker for identification of unknown animal species. L1-L5 lanes refer cow, camel, goat, sheep & chicken with 307, 385, 787, 763 & 470bp respectively, L7-L8 lanes refers cat & dog with 700 & 370bp respectively, similarly, L11-L15 lanes refers donkey, duck, rabbit, horse & human with 600, 500, 750, 800 & 300bp respectively. While L6, L10 and L16 lanes represents100kb DNA ladder.

**Figure 1:**
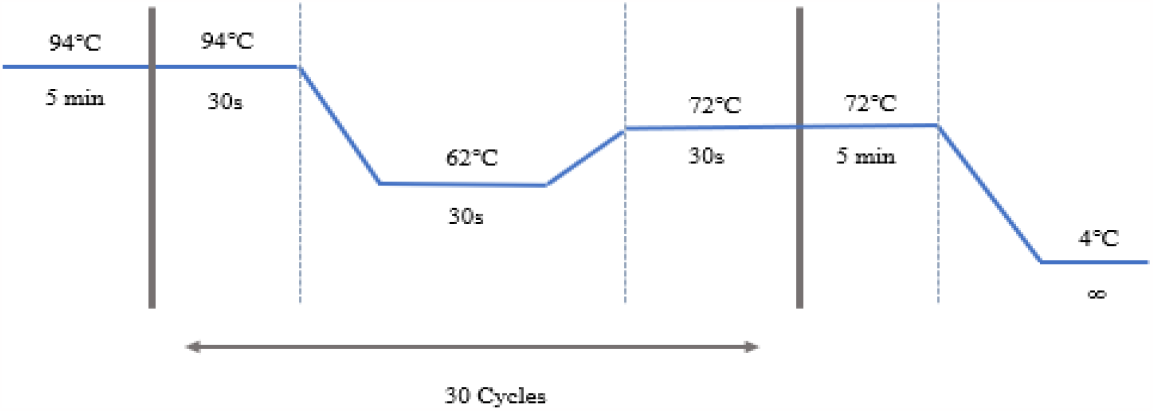
Thermal cyclic conditions for PCR

**Figure 2:**
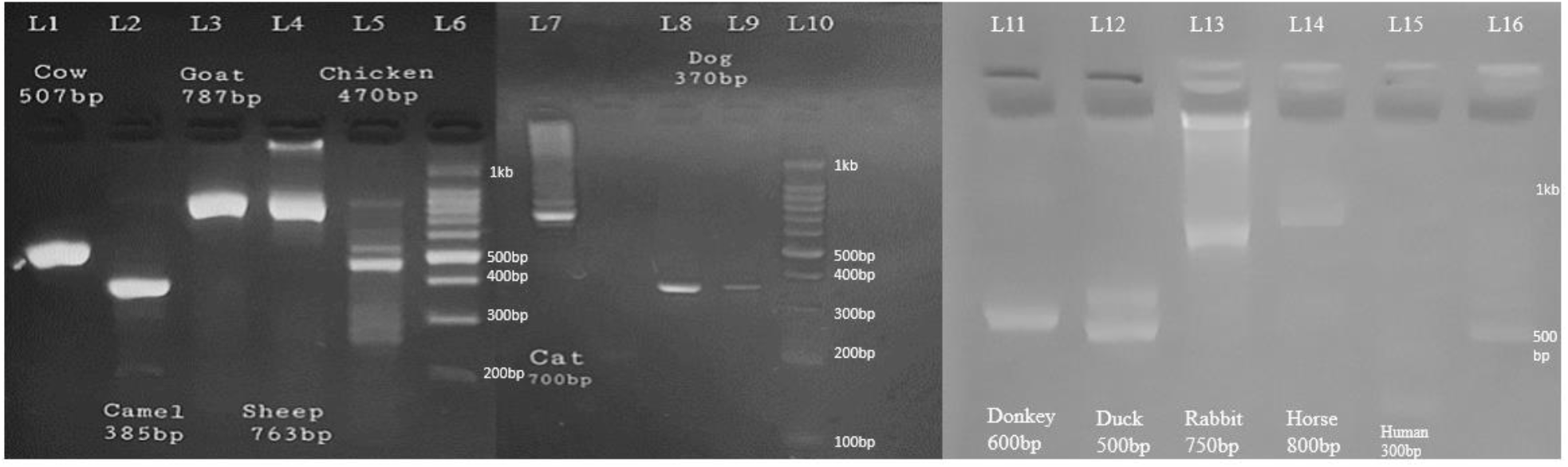
Gel electrophoresis for PCR products amplified with universal primers.

The amplified amplicon is not same in the subject species as evident from the figure 2. So, the selected mitochondrial DNA marker can be used for species identification. Further, validation was also performed using the same set of primer from the in-silico PCR tool from UCSC genome browser (https://genome.ucsc.edu/cgi-bin/hgPcr).

## Discussion

The primary objective of the current study is to discern and categorize the diverse animal species meat present in food products. In response to an escalating awareness among consumers regarding food safety and security, researchers are compelled to enhance the efficacy of existing techniques and develop more robust methodologies to detect potential adulterations in meat and food items. Among the current array of techniques, the PCR stands out as exceptionally efficient, making it the optimal choice for species identification. This preference is attributable to its cost-effectiveness, sensitivity and precision [8]. This technique has demonstrated significant success in identifying potentially adulterated meat and meat products.

Matsunaga et. al employed a multiplex PCR approach to identity the six distinct meat species [9]. Park Y.H et. al. utilized a species-specific multiplex PCR a comprehensive identification was achieved through the implementation of 12S RNA and mitochondrial *Cytb* gene to identify eight meat species [10]. The development of a singular set of universal oligonucleotides targeting the 16S rRNA mitochondrial genes enables the identification of traces of multiple animal species in food products through a single PCR reaction. This direct approach to species identification for ensuring food authenticity has evolved into a more precise and specific methodology. The selected animals exhibit conserved sequences with subtle variations, as evidenced by their cladogram depicting a common root with two main clades. The upper clade encompasses four nodes, incorporating chicken, duck, cat, camel, and donkey, while the lower clade features six nodes, encompassing seven animal species: human, rabbit, cow, goat, sheep, dog, and horse (figure 3).

**Figure 3:**
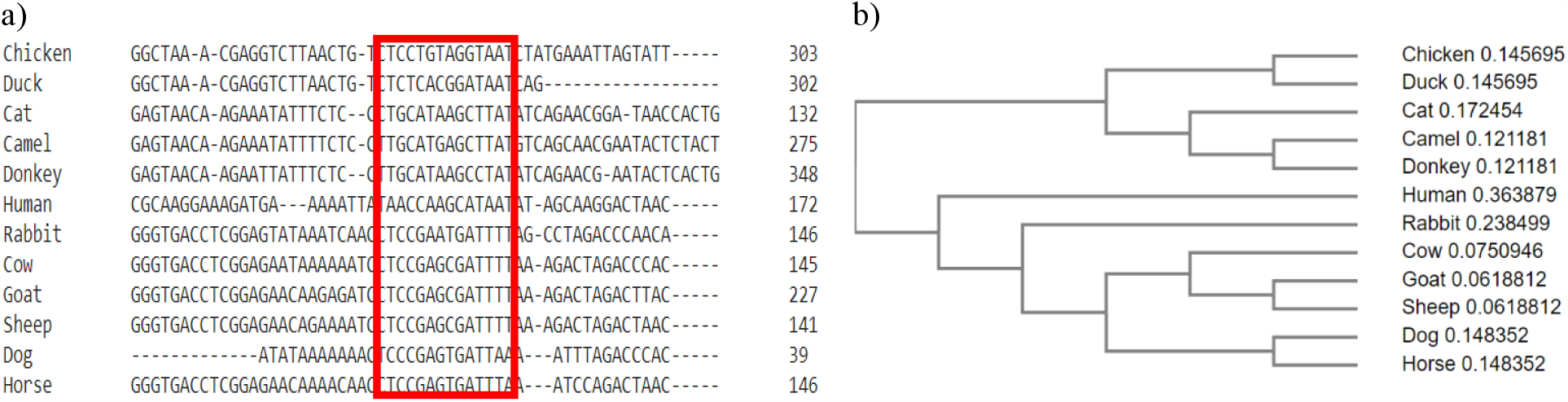
a) Multiple sequence alignment of the consensus sequence of 16S rRNA in twelve animal species and b) cladogram showing relatedness of the subject animals on the basis of selected gene

## Conclusion

Using universal primers designed for the 16S rRNA gene, this study effectively distinguishes twelve indigenous domestic animal species in Pakistan. The amplified targets show species-specific traits, preventing similar results among different species. These findings provide a crucial basis for identifying unknown meat species, serving as a robust database for comparisons in forensic and food authenticity assessments, thereby enhancing accuracy.

## Notes

### Competing Interest Statement

The authors have declared no competing interest.

